# Qualitative modeling of signaling networks in replicative senescence by selecting optimal node and arc sets

**DOI:** 10.1101/389692

**Authors:** Theodore Sakellaropoulos, Ilona Binenbaum, Maria Lefaki, Aristotelis Chatziioannou, Niki Chondrogianni, Leonidas G. Alexopoulos

**Affiliations:** School of Mechanical Engineering, National Technical University of Athens, Athens, Greece; Institute of Biology, Medicinal Chemistry and Biotechnology, National Hellenic Research Foundation; Department of Biology, University of Patras

## Abstract

Signaling networks are an important tool of modern systems biology and drug development. Here, we present a new methodology to qualitatively model signaling networks by combining experimental data and prior knowledge about protein connectivity. Unlike other methods, our approach does not focus solely on selecting which reactions are involved but also on whether a protein is present. This allows the user to model more complicated experiments and incorporate more knowledge into the model. To demonstrate the capabilities of our method we compared the signaling networks of young and replicative senescent human primary HFL-1 fibroblasts, whose differences are expected to be due mainly to changes in the expression of the proteins rather than the reactions involved. The resulting networks indicate that, compared to young cells, aged cells are not as responsive to insulin stimulation and activate pathways that establish and maintain senescence.

**Author summary:** Cells have developed a complex network of biochemical reactions to monitor their environment and react to changes. Although multiple pathways, tuned to identify specific stimuli, have been discovered, it is generally understood that the signaling process typically involves multiple pathways and is context depended. Consequently, reconstructing the signaling network utilized by cells at any given moment is not a trivial task. In this article, we report on a novel logic-based method for identifying signaling network by combining experimental data with prior knowledge about the connectivity of the involved proteins. Unlike other methods proposed so far, our method uses data to evaluate the presence or absence of reactions and proteins alike. We reconstructed and compared the signaling network of human primary HFL-1 fibroblasts as they undergo replicative senescence in the presence of 6 different stimuli. The resulting networks indicate that, compared to young cells, senescent cells are not responsive to insulin stimulation and activate pathways that are known to establish and maintain senescence.

## Introduction

A central challenge of Systems Biology is to develop mechanistic models that explain cell behavior as illustrated by high-throughput experimental data. Models describing cell signaling are of particular importance in understanding how a cell interacts with its environment and have been widely used in drug discovery [1,2] since many drugs target the signaling process by inhibiting specific interactions [3]. A major challenge in dealing with signaling networks is that there are no golden standards [4] as they are context dependent. Information about interactions is scattered among many databases which have different policies with respect to what constitutes an interaction, which interactions are included and the format to encode them [5,6]. Moreover, several researchers have pointed out that there is a low consensus among these databases, presumably due to the low specificity of the currently available techniques to identify interactions [7–9]. Thus, the currently available interactome serves more as a palette of possible interactions rather than a snapshot of what is going on inside a cell.

As a result, many algorithms have been proposed for constructing signaling networks by adapting the interactome (prior-knowledge about connectivity) to experimental data in order to produce a network specific to the mechanism under investigation [10]. Experimental data are usually derived from perturbation experiments aiming to highlight the dependencies between some nodes of the network. There are two main steps in designing these algorithms; first, a mathematical formalism is selected to interpret the signal transduction mechanism and second an optimization process is carried out to tune the parameters of the model according to the data. Deciding the appropriate formalism depends primarily on the questions being asked and the type of data available [11]. Some approaches consider the prior knowledge network fixed and use data to tune its dynamic behavior while others use data to adjust the connectivity. Methods belonging to the latter category are more suitable for dealing with large signaling networks due to the limitations described in the previous paragraph. Once a formalism is selected, a network can be used to simulate the signaling process and thus make predictions about the involved entities. This predictive ability allows one to access the quality of the network, and ultimately modify it to improve it, by comparing its predictions with experimentally observed data.

To the best of our knowledge, the methods proposed thus far in order to reconstruct signaling networks focus on selecting a subset of the known reactions that can optimally simulate the observed behavior. This approach is justified for two main reasons. First, as discussed above, a lot of the reported reactions are considered to be false positives or context dependent. Second, cells can be probed in the presence of inhibitors, like drugs or small molecules, that inhibit specific reactions and thus the effect of these interactions can be evaluated directly. However, signaling networks can also be altered in cells by the responsiveness of the involved proteins, either because they are saturated, the cell under-expresses them or researchers have knocked them down, using CRISPR or shRNA for example.

In this paper, we propose a novel logic-based method for identifying signaling networks using signed directed graphs, also known as interaction graphs (IG) [12]. Interaction graphs provide a description powerful enough to capture global dependencies and make qualitative predictions so they fall under the second category of data-driven methods. We use mixed integer programming (MIP) to optimize the networks which, despite being NP-hard in principle, have been proven effective in dealing even with large instances of these problems [13–15]. Our method expands upon our previous work [16] on the subject by expanding the scope of the optimization to include both reactions/arcs and proteins/nodes of the network. As a case study, we reconstructed the signaling network of human primary fetal lung fibroblasts (HFL-1) as they undergo replicative senescence. In the case of aging, we suspect that the reactions involved in the investigated processes should remain unchanged over time and only the expression or responsiveness of the involved entities should be affected.

## Materials and methods

### Preliminaries

We assume that we are given a signed directed graph *G* = (*N, A, σ*) representing our prior knowledge about the connectivity of the entities involved. Sign directed graphs, also known as interaction graphs (IG), extend the notion of directed graphs by assigning a sign to every arc *σ : A* → {−1, +1}. In our model, the nodes (*N*) of an IG can assume one of three possible states: up-regulated (+1), down-regulated (−1) and unperturbed/basal state (0). The sign of an arc represents the effect or influence the parent-node exerts on the child-node (see sign consistency later). To allow for more complicated scenarios to be modeled, each node and arc is associated with a—possibly empty—set of inhibitors *I_n_* and I_α_ respectively. These may correspond to actual inhibitors, like drugs and small molecules which can inhibit particular reactions or other types of inhibition such as shRNA or CRISPR which can knock-down nodes completely. Inhibitors function as switches that force their associated elements to their basal state when activated. To keep the problem description concise we define the inhibitor sets as *I_n_, I_a_* ⊆ *N* × {−1,1}, ie that they can be described by the state of some nodes of the graph.

For our case study, the MetaCore database was used to construct the prior knowledge network (PKN). MetaCore is a manually curated database of signaling networks that includes both positive and negative interactions as well as quantitative metrics of confidence for each interaction like the number of publications supporting it. We curated the network in order to include only observable and controllable nodes, i.e. nodes that are connected to both signals and stimuli. The final PKN consisted of 90 nodes and 262 interactions (Fig 1). Finally, we used the number of references to prioritize which reactions would be included in the final network.

**Fig 1.**
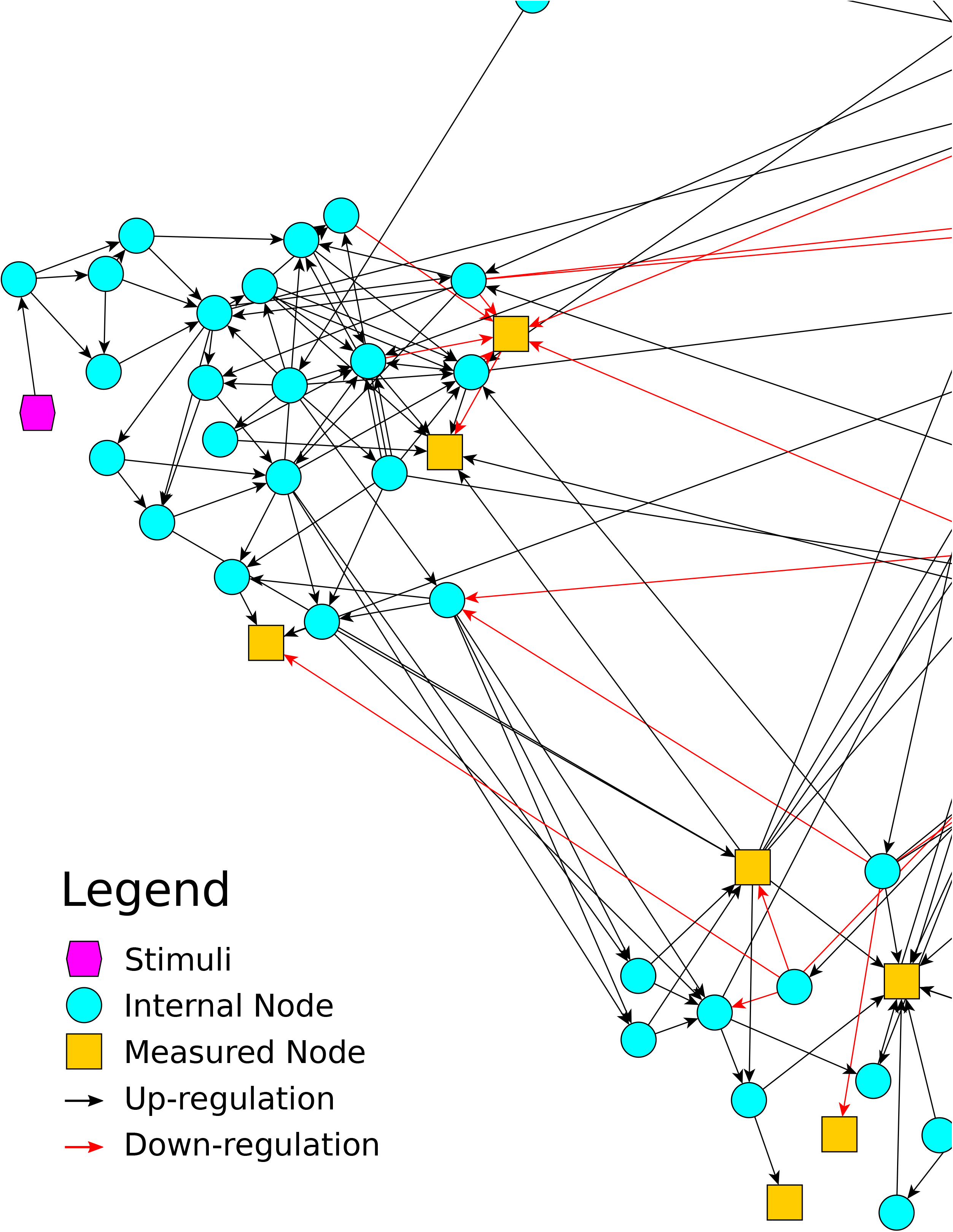
Prior Knowledge Network. Constructed by manually curating Metacore. It contains 90 nodes and 262 edges.

Apart from prior knowledge about the connectivity, our method also requires a set of experimentally observed dependencies in order to refine the PKN so as to simulate them as closely as possible. Dependencies can be observed via perturbation experiments where some nodes of the network are perturbed (set in a non-basal state) and the state of some other nodes is recorded. In the context of signaling networks, these experiments correspond to measuring the phosphorylation status of intra-cellular proteins upon perturbing the cells. We assume that these experiments capture a *steady-state shift.*

Experiments are modeled as a set *E* of tuples. Each tuple *e ∈ E* consists of two functions; a perturbation function *p_e_* : *N* → { — 1,0,1} indicating which nodes are initially perturbed and *m_e_* : *M_e_* → [−1,1] indicating the state of the measured nodes in experiment *e M_e_* ⊆ *V*. A perturbation of zero indicates an unperturbed node, to force a node to assume the state zero an inhibitor for the corresponding node must be added. On the other hand, a measurement of zero is not equivalent to no measurement, that is why *m_e_* are not defined over all nodes *N*. Another important distinction between *p_e_* and *m_e_* is that the first maps to discrete values indicating that perturbations are known with certainty while measurements are not. The image of *m_e_* is a “fuzzy” relaxation of the three distinct states where the distance from each discrete value {−1, 0,1} is inversely proportional to the modeler’s belief that the node measured was in the corresponding state.

To simulate the outcome of a perturbation experiment, we must produce a *sign consistent labeling* of the nodes. The notion of sign consistency is discussed in detail in [17]. Here, we briefly outline the aspects that we are going to use to simulate the signaling process. A sign consistent labeling *l* : *N* → {-1, 0,1} abides by a set of rules that restrict the possible state-combinations of adjacent nodes. First, every perturbed node must be labeled with the intended sign

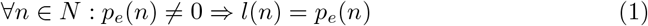

Second, the state of every non-perturbed node must be equal to the state of at least one of its predecessors multiplied by the sign of the connecting arc

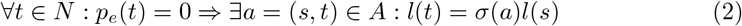

Notice that this rule produces a “weakly consistent” network.

### ILP Formulation

In this section, we describe a mixed integer program to prune the original PKN in order to identify a subnetwork that can be labeled sign consistently in a way that minimizes the divergence between labels and measured states. The main idea of the formulation is to use the constraints in order to force a sign consistent labeling and then use decision (binary) variables to select nodes and arcs in order to minimize the objective function. We begin by presenting the indices and the variables utilized together with their interpretation and then proceed to describe the constraints and finally the objective function of the problem.

Variables are indexed by *n ∈ N* to indicate the node, *a ∈ A* to indicate the arc and *e ∈ E* to indicate the experiment. For each node *n*, a binary variable *v_n_* is introduced to model whether it is part of the optimal sub-network. To model a node’s state in experiment e, two binary variables are introduced 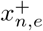 and 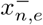 modeling whether the it is up or down regulated respectively. For each arc *a*, a binary decision variable *y_a_* is introduced to model whether it belongs to the optimal sub-network and two continuous variables 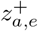 and 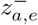 to model whether, in experiment *e*, the signal originated from an up or a down regulated node. The asymmetry in the *x* and *z* variables, the former being a decision variable while the latter not, is meant to accommodate the degrees of freedom that sign consistency allows. In particular, the fact that each node has to be consistent with *at least* one of each predecessor allows the optimizer to select the label that would result in a better overall fit. The relationship between the x and z variables is summarized visually in 2.

The constraints modeling sign consistency can be grouped into two main groups, those regulating the state of the nodes and those regulating the state of the arcs. For notational convenience, we use the symbols *a_s_* and *a_t_* to indicate the source and target node of an arc *a* = *s → t* and ± to indicate “double” constraints both for the positive and negative variables involved.

We begin with the arc regulating constraints. For every arc *a ∈ A* in every experiment *e ∈ E* the following constraints must hold:

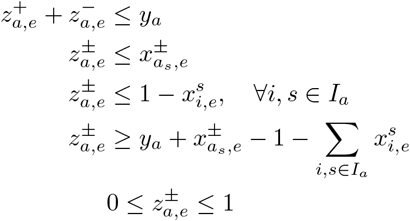

The first four constraints implement an AND gate of necessary and sufficient conditions for an arc *a* to transduce a signal in experiment *e*. The three conditions are; first, the arc must be included in the final sub-network, second, the source node must be in the appropriate state and third no inhibitors should be active. If all of these conditions are satisfied then the fourth, constraint ensures that the arc will become active. The first and the last constraint also ensure that *z*^+^ and *z*^−^ behave as mutually exclusive binary variables. The reason we do not have to explicitly designate z as binary is that their value is uniquely determined for every combination of the actual decision variables of the problem.

For every node *n ∈ N* and every experiment *e ∈ E* the following constraints must hold:

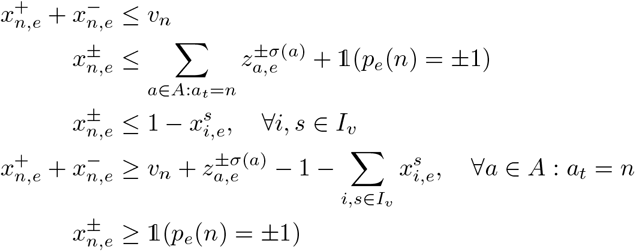

where 1 is the identity function—equals 1 when the condition is met.

Again, the first four constraints implement an AND gate of necessary and sufficient conditions for a node to be up or down regulated. These conditions are; first, the node must be included in the final sub-network, second, unless it’s perturbed, at least one reaction must activate it and third, no inhibitor should be active. If all of the three requirements are met then the fourth constraint will force the node to assume a non-basal state. The first constraint also enforces mutual exclusivity between the two active states of a node, as it cannot be both up and down regulated. The fifth constraint guarantees that the perturbed nodes have the intended state and initiates the signaling process.

The sum at the left-hand side of the fourth constraint allows the optimizer an extra degree of freedom in labeling a node. In particular, unless forced by another constraint, a node can become either up or down regulated regardless of the sign of *z*, and the optimizer has to decide the best option. If there are no conflicting influences, then there is no decision to be made since the second constraint would have already forced one of the two *x* variables to zero. Because of this constraint, *xs* have to be binary variables, unlike the *zs*.

The final subnetwork will always be an acyclic graph rooted at the perturbed nodes and branching towards the measured nodes. Nonetheless, the prior knowledge network may contain loops. Loops present a challenge for logic-based methods since they can be self-activated without external perturbation. Because the up and down regulated states are mutually exclusive, negative feedback loops are automatically handled by the constraints presented thus far. However, for the case of positive feedback loops we have to include an extra set of constraints and variables. In particular, for every node *n ∈ N* we introduce a variable *d_n_* and the following constraints:

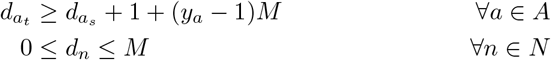

In the presence of a cycle, these constraints will push the *d* variables involved to grow without bound and thus by bounding them up by *M* we force the optimizer to break the cycle by removing some arcs. *M* represents a “big” constant and can be thought of as the longest distance between any pair of nodes and thus the best “uninformative” value is *M* = |*N*| – 1, corresponding to a path spanning all the nodes. It’s also worth mentioning that every node will have a larger *d* variable than all its predecessors and thus the order of the *d* variables corresponds to a topological ordering of the nodes which is only defined for acyclic graphs.

To produce the subgraph that best simulates the observed cell behavior, we set up least-squares objective function to minimize the difference between the observed and predicted response. In more detail, if 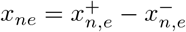 is the predicted state of node n in experiment e then the objective is to minimize (*x_n,e_*,– *m_e_*(*n*))^2^ across all experiments, where mò is the measured state of the node as discussed above. This functional is equivalent to the following linear function:

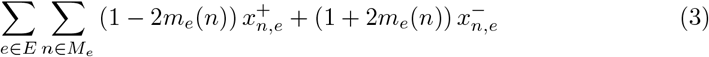

Sparsity incentives were also encoded in the objective function in the form of weights over the variables *v* and *y*. The weights were derived from the number of citation supporting each reaction and were normalized in the [0, −1] interval where −1 was assigned to the reaction with the most citations. In this way, the trade-off between sparsity and accuracy was 1:4.

For our case study of aging, we reckoned that during aging *in vitro* cells only alter the expression of the involved proteins rather than their signaling pathways, thus we constrained the networks of young and old cells to share the same set of reactions. The two networks were allowed to differ only with respect to the set of active nodes. These constraints were facilitated by “coupling” the y variables of the two networks while allowing the v variables to differ. In this manner, both sets of experiments contribute in determining the reactions involved.

Finally, because of the discrete nature of the problem, many solution can be optimal or nearly optimal. To address this issue, we generated multiple near optimal solutions and for each variable we averaged its value across all weighted by the value of the objective function.

### Experiments

We maintained human primary fetal lung fibroblasts (HFL-1) in culture from a young stage until they reached a terminally senescent (old) stage after 50 replication cycles. In the young stage the cells were replicating every 24 hours while in the old stage they had not replicated for a month. Cells were seeded in 96-well plates at 20000 cells/well for 24 hours prior to stimulation. After 24 hours cells were stimulated by adding the stimulants to the cell medium at a concentration calculated to reach the target concentration in the cell supernatant. Cell lysates were collected at 5 and 25 minutes following the cytokine stimulation. The 5 and 25 minute time points were identified in a preliminary experiment as the optimal time points to capture early phosphorylation activities. A panel of 18 phosphoproteins upon stimulation with 6 cytokines was used to interrogate the cells. The panel was selected to represent known proliferation and senescence regulating pathways. Stimulant concentrations were selected after pre-screening experiments. The cytokines used to stimulate the cells and their target concentrations were: *EGF* (100 ng/ml), *IL1A* (50 ng/ml), *TGFA* (200 ng/ml), *TNF* (100 ng/ml), *IGF1* (200 ng/ml) and *INS* (1000 ng/ml). The interrogated phosphoproteins were: *AKT1, CREB1, PTK2, GSK3A, HSPB1, NFKBIA, JUN, MAPK12, MAPK3, MAPK9, MAP2K1, TP53, PTPN11, STAT1, STAT3, STAT5, STAT6* and *RELA.* Basal protein phosphorylation was measured in absence of any stimulant in DME cell medium.

HFL-1 human primary embryonic lung fibroblasts were purchased from ECACC (ECACC 89071902). Cells were cultured in Dulbecco’s modified Eagle’s (DME) medium (Invitrogen, Carlsbad, CA, USA) supplemented with 10% fetal bovine serum (Invitrogen), 100 units/mL penicillin, 100 μg/mL streptomycin, and 2 mM glutamine (complete medium) and maintained in a Binder Incubator at 37°C, 95% air, 5% CO_2_. HFL-1 fibroblasts were seeded at a density of 2 × 10^5^ cells per 75 cm^2^ flask, were subcultured at a split ratio of 1:2 when cells reached confluence, until they entered senescence at about 50 CPD, as described before [18]. In all experimental procedures described below early passage (young; CPD <25) and late passage (senescent CPD >50) HFL-1 cultures were used. Cells were fed approximately 16 hours prior to the assay and cell number were determined in duplicates using Coulter Z2 counter (Beckman Coulter, Nyon Switzerland). The *IGF1* (100-11), *EGF* (AF-100-15), *IL1A* (200-01A) *TGFA* (100-16A) and *TNF* (300-01A) cytokines used for stimulation were purchased from Peprotech (PeproTech, NJ, USA), and INS (INS I9278) was obtained from Sigma-Aldrich (Sigma-Aldrich, Shanghai, P.R. China). Measurements were carried out in 96-well plates using Luminex’s xMAP technology with custom assays by ProtATonce Ltd.

The median fluorescent intensity was used to summarize the results of the Luminex measurements. To map values to the [−1,1] interval required by the algorithm, fold changes were computed between the cytokine-stimulated state and the basal state (DME) and passed through the Gaussian error function. The two time points (5 and 25 minutes) were aggregated into a single “early” time point by taking the maximum absolute value which is analogous to an OR gate. See S1 Supporting Information for more details about the data set.

Technical variability for bead-based phosphorylation assays has been studied before in [19], where they estimated the coefficient of variance due to the experimental protocol and pipetting errors to average in the range of 5–15%. To access the impact of such variability in our results, we generated artificial datasets by adding noise to our MFI readouts and rerun the analysis. We repeated this processes 10 times for three different levels of noise 5, 10 and 15%, and computed the Jaccard similarity index of the resulting network with the original network. We observed that a CV of 5% has virtually no effect in the results while for larger values the networks start to diverge. The results are summarized in Figure 3.

**Fig 2.**
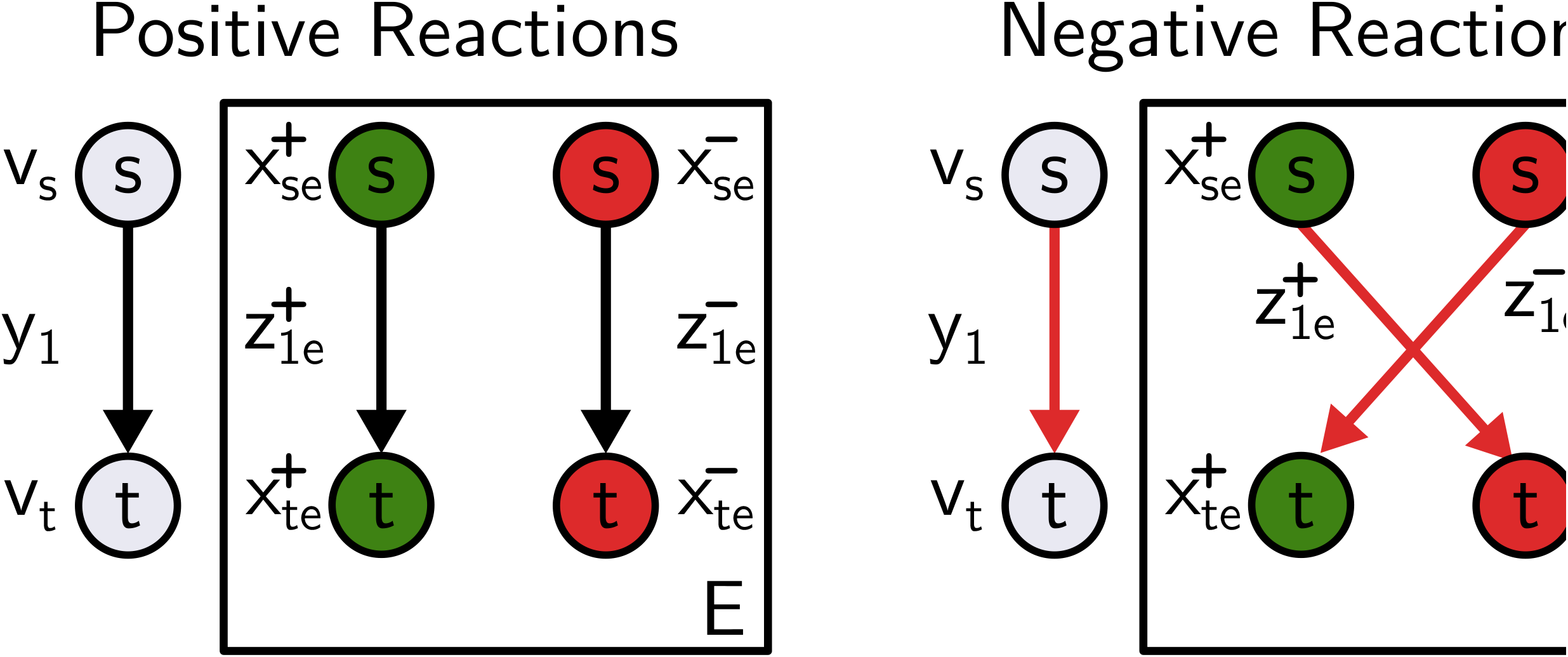
Arc decomposition. Positive and negative effect are modeled by different variables and the dependencies between them characterize the reaction.

**Fig 3.**
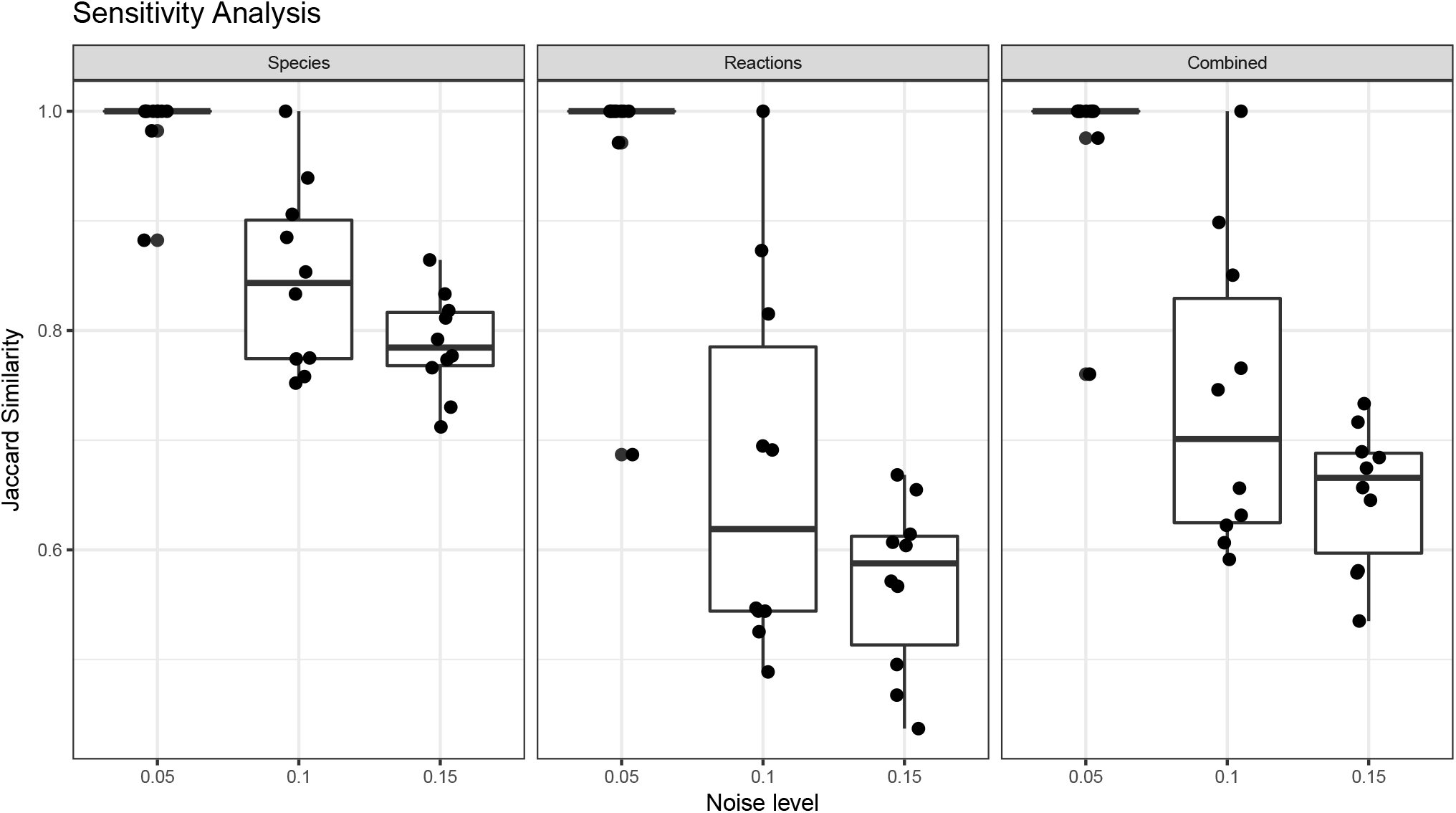
Sensitivity Analysis. 10 randomized datasets were generated for each noise level and the optimal networks were reconstructed and compared with the original dataset. The comparison was based on the presence of reactions (*y*), species (*v*) or both.

## Results

Figure 4 summarizes the measured states *m_e_* of every phosphoprotein measured upon stimulation with every cytokine both for young and senescent cells. As expected, young cells appear to be much more responsive across the board. We observed that the signaling proteins that are involved in proliferation including AKT Serine/Threonine Kinase 1 *(AKT1*), *cAMP* Responsive Element Binding Protein 1 *(CREB1*), Mitogen-Activated Protein Kinase Kinase 1 *(MAP2K1*), Mitogen-Activated Protein Kinases 3 and 9 *(MAPK3, MAPK9*), Protein Tyrosine Kinase 2 *(PTK2*), Protein Tyrosine Phosphatase, Non-Receptor Type 11 *(PTPN11*) and Signal Transducer and Activator of Transcription (STAT) family members 1, 3 and 5 *(STAT1, STAT3, STAT5*) are substantially more activated in young than senescent cells. Senescent cells show a higher activation of *STAT6* when stimulated by *EGF* or *IL1A.* Stimulation with *EGF* also provides a higher response in senescent than young cells in the *NFKB* Inhibitor Alpha *(NFKBIA), RELA, STAT3, STAT6* and Tumor Protein P53 *(TP53*) and overall it seems that *EGF* is the growth factor that senescent cells are most responsive to. Notably, Glycogen Synthase Kinase 3 Alpha *(GSK3A), TP53, STAT1, STAT5* and *MAPK9* respond in opposite ways in young and senescent cells across multiple stimulants. Heat Shock Protein Family B Member 1 *(HSPB1*) and Jun Proto-Oncogene *(JUN*) are more responsive in young cells as well.

**Fig 4.**
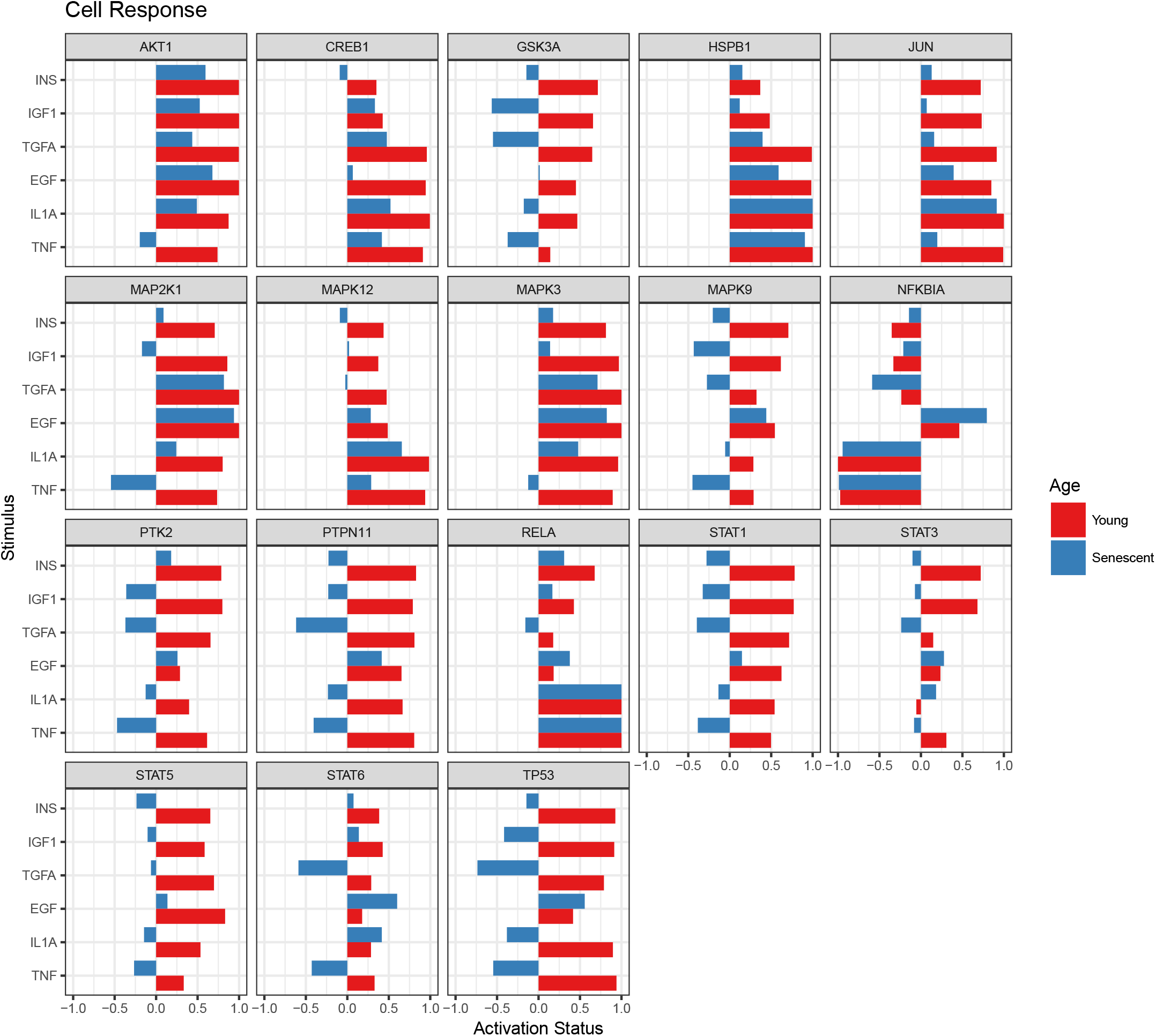
Cell Response to different stimuli. Panels correspond to measure phosphoproteins. On the x-axis is their measured status (−1: down-regulated, 1: up-regulated) and on the y-axis the stimuli used in every experiment.

To gain a more mechanistic insight into what drives this change in response, we run our algorithm for young cells, replicating every 24 hours, and old cells that have entered the senescence state. The resulting network is shown in Fig 5. The predicted status of the signals is shown in Fig 6.

**Fig 5.**
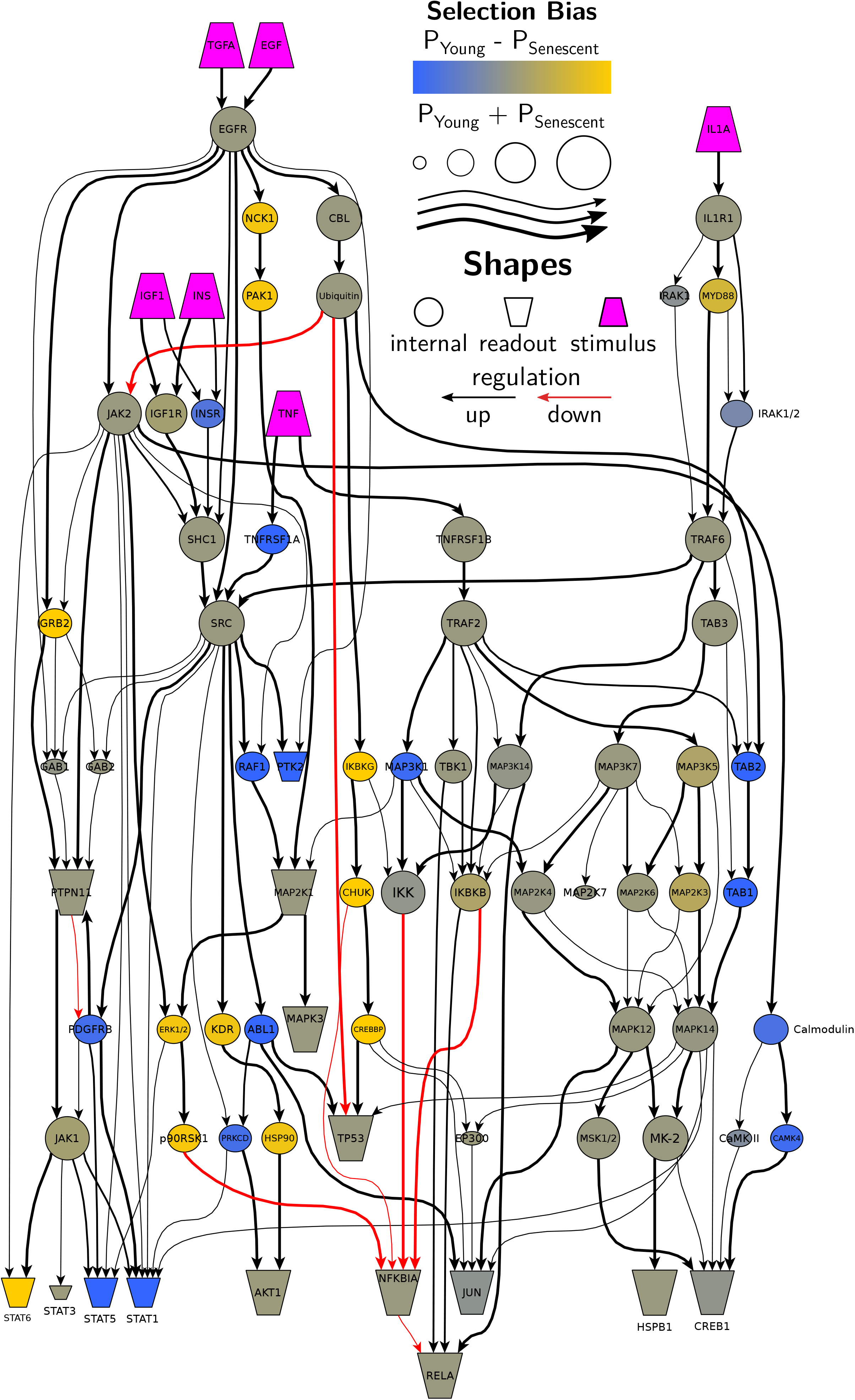
Network Comparison Average network produced. The color hue corresponds to bias towards young or senescent cells. Node and arc size corresponds to probability of inclusion in a solution. The shape of the nodes is used to distinguish between signals, stimuli and internal nodes while the color of the arcs between up and down regulating reactions.

**Fig 6.**
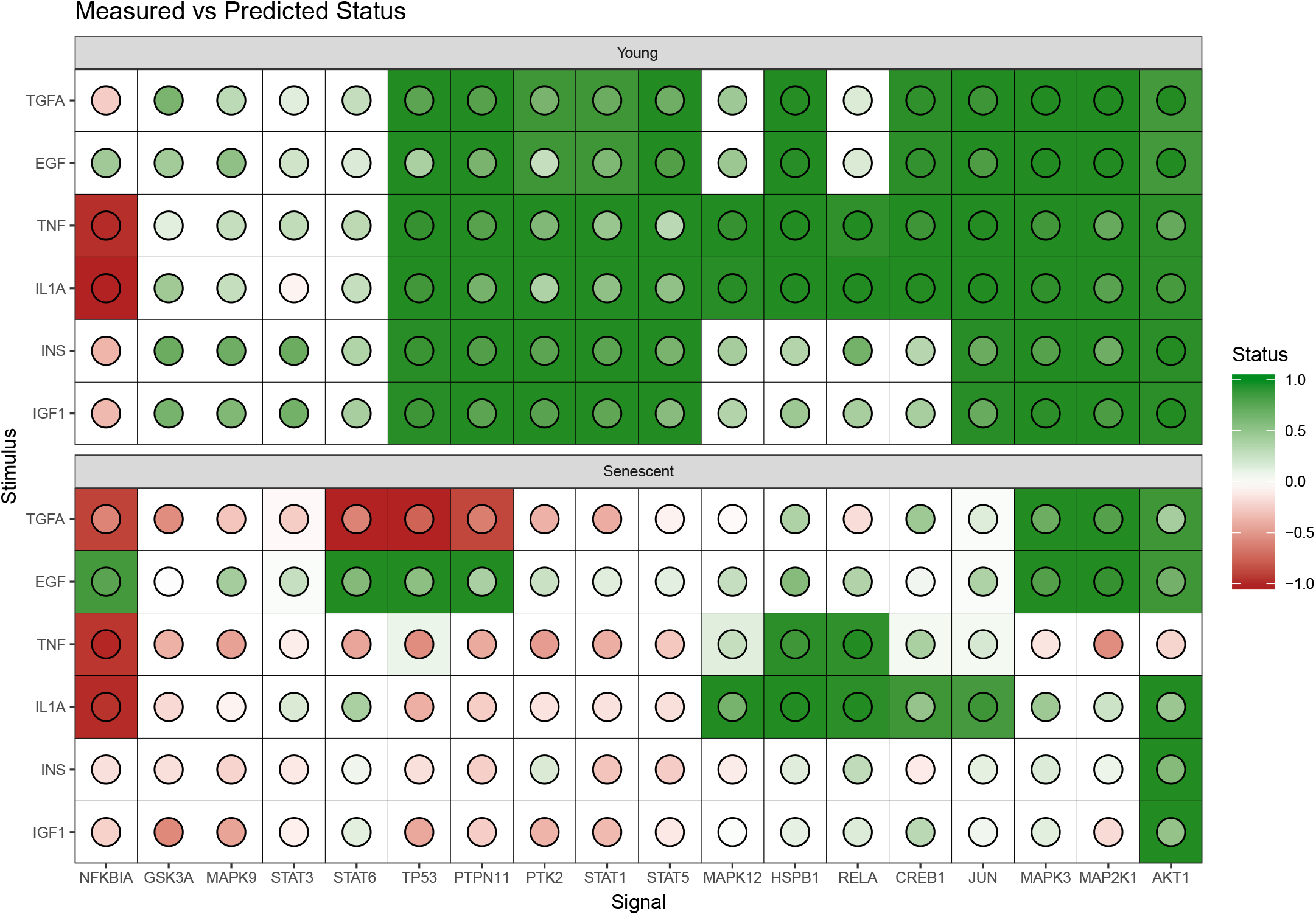
Network Predictions. Every tile corresponds to a stimulus-signal combination. The tile color corresponds to the predicted status of the signal while the color of the enclosed disk corresponds to the measured status. The predicted status is averaged across multiple solution to rendered continuous.

The basal protein levels of several of the interrogated proteins were revealed in young and terminally senescent cells. We revealed differences in the basal expression of several of them Fig 7.Basal protein phosphorylation was measured in absence of any stimulant in DME cell medium showing that phosphorylated forms of the indicated protein were also existing in the cells.

**Fig 7.**
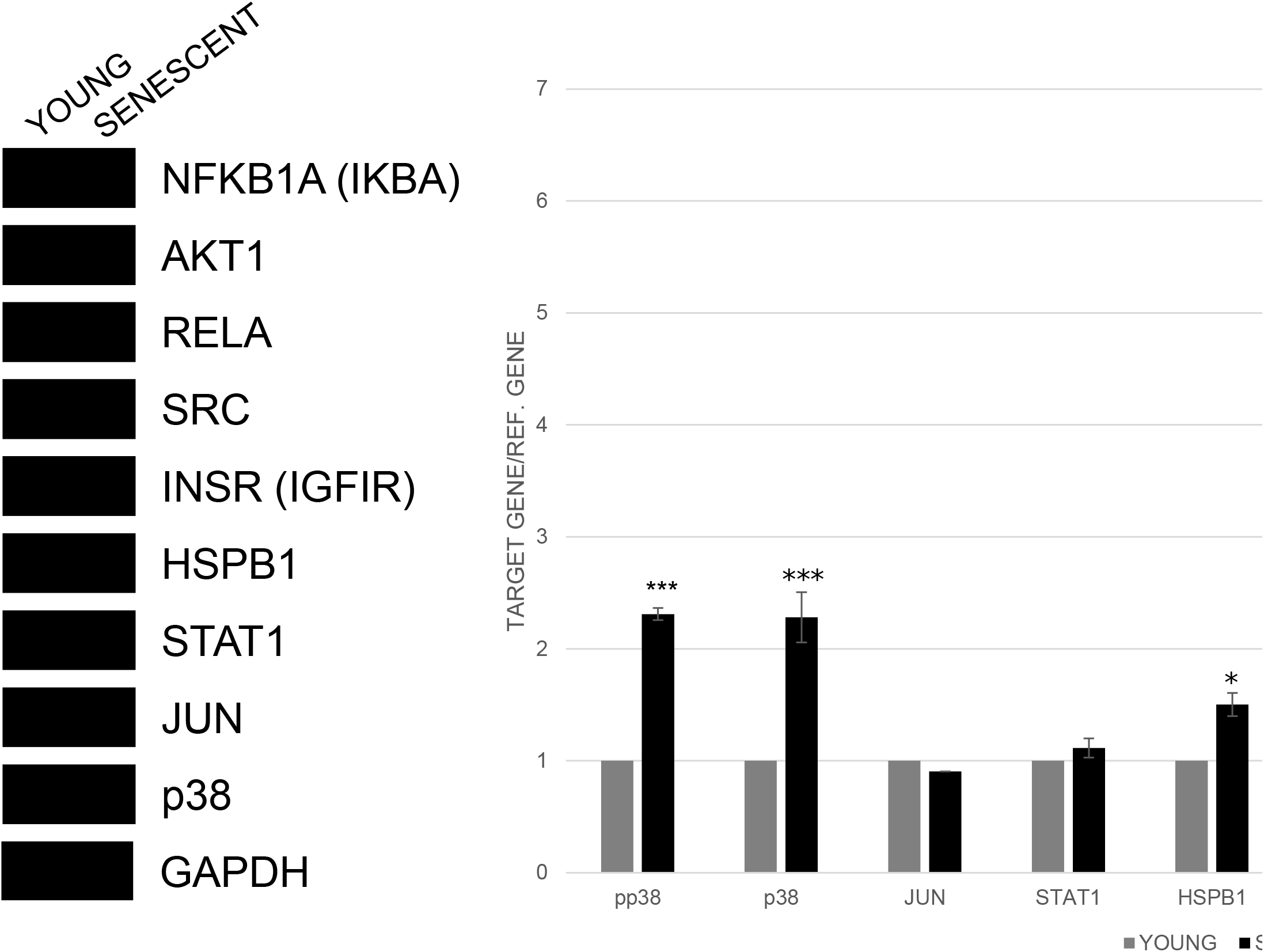
Basal protein levels immunoblot. Immunoblot analysis and the relative quantification in young and terminally senescent HFL-1 human primary fibroblasts. GAPDH was used as loading control. Error bars denote ± SEM. * P < 0.05, ** P <0.01, *** P <0.001

In our model, we observe that common but also different phosphoproteins can be utilised by young and senescent cells. The insulin-like growth factor 1 receptor *(IGF1R)* is predicted to be more responsive to stimulation both with the insulin-like growth factor 1 *(IGF1*) and insulin *(INS*) than the insulin receptor (INSR), in both types of cells. However, young cells appear to utilise *INSR* more often than senescent cells.

Senescent cells appear to have more downstream pathways activated in response to *TGFA* and *EGF.* While the Epidermal Growth Factor Receptor *(EGFR)* is activated in both types of cells, leading to predicted activation of Cbl Proto-Oncogene *(CBL)* and Janus Kinase 2 *(JAK2*), in senescent cells NCK Adaptor Protein 1 *(NCK1*) and Growth Factor Receptor Bound Protein 2 *(GRB2*) show a higher probability of being activated as well, leading to higher probability of downstream activation of P21 (RAC1) Activated Kinase 1 *(PAK1*), *STAT6* and *ERK1/2. STAT6* has been found to be involved in the induction of cellular senescence by IL-4 [23]. Moreover, in senescent cells *NFKBIA,* is predicted to be more often down-regulated by upstream inhibitors, Conserved Helix-Loop-Helix Ubiquitous Kinase *(CHUK*) and Inhibitor Of Nuclear Factor Kappa B Kinase Subunit Beta *(IKBKB*) and *RelA/p65* is slightly more likely to be up-regulated. These signaling events can lead to the activation of *NFKB* signaling that establishes and maintains cellular senescence [22]. Additionally, signaling through the *TNF* receptor 1 *(TNFRSF1A)* that leads to downstream activation of *SRC,* is also predicted to happen predominantly in young cells, while the *TNF* receptor 2 *(TNFRSF1B*) is equally responsive in both types of cells.

On the other hand, in young cells we predict higher probability of activation of proteins involved in cell growth and proliferation such as *STAT1, STAT5,* and *PTK2,* which is confirmed for all three by our experimental data. The basal phosphorylation levels of these proteins are also significantly higher in young cells. In response to *IL1A, MYD88* is more likely to be phosphorylated in senescent cells while IL1 receptor activated kinases *(IRAK1/2*) are more likely to be responsive in young cells. *MYD88* activation through the interleukin receptor, followed by recruitment of IRAKs has been associated with the activation of the senescence-associated secretory phenotype *(SASP*) [29].

## Discussion

In this article, we present a novel framework for reconstructing signaling networks using phosphoproteomic data and prior knowledge about their connectivity. The framework presented expands our previous work on the subject by allowing species and reactions to be modeled independently. This decomposition renders the framework more flexible to deal with evolving systems, like aging and senescence, and also allows the user to incorporate more information into the model both with respect to species and reactions. For example, cells can be interrogated in the presence of drugs or small molecules that inhibit specific interaction but also upon knocking down specific genes and thus removing their respective proteins.

For the case study presented, the reaction set was constrained to be the same across both populations while the node set was allowed to vary but the opposite configuration could have been implemented just as easily. The configuration used allowed networks to differ only due to changes in the responsiveness levels of the proteins while also allowing more data points to inform which reactions should be included. More generally, within the presented framework, one can impose soft or hard constraints on nodes and/or reactions without necessarily compromising the accuracy of the model, since the two mechanisms allow the optimizer to work around them if necessary. From the sensitivity analysis presented, we observed that, despite the fact that reactions were constrained to be the same for the 2 populations, they were more sensitive to noise. This is due to their relatively moderate effect on the overall connectivity compared to the effect of removing a node from the network. Thus by extending the scope of optimization to species as well as reactions we render our model less sensitive to overfitting.

Similarly to our previous work, the proposed formulation can handle both positive and negative interactions as well as cycles in the prior knowledge network transparently. However, the resulting networks are acyclic so it cannot model dynamical behavior within a network. Also, as with all logic-based approaches, our framework lacks the ability to describe the kinetic aspects of the network [11]. Because of the MIP implementation, the networks constructed are optimal with respect to the proposed framework or at least within a pre-specified distance from optimal [21]. Moreover, modern MIP libraries like CPLEX and Gurobi, allow the enumeration of multiple Pareto-optimal or near optimal, solution. We run our method on synthetic data, that respected sign consistency, and were able to reconstruct the generated signatures in every occasion (data not shown).

The number of variables/constraints required is *O*(*2k*(*N* + *A*)) where *N, A* is the number of species and interactions in the PKN and *k* is the number of experiments simulated. The 2 is due to the fact that we need to distinguish between up and down regulation. So depending on the infrastructure and the number of experiments it can handle networks for thousands of species. In our experience, the bottleneck is the number of experiments k. Furthermore, from a computational point of view, the number of proteins is typically only a fraction of the number of reactions resulting in a possibly significant shrinkage of the search space, if the optimization is carried over the proteins only.

To demonstrate our framework, we modeled how cells alter their signaling network as they senesce *in vitro*. In particular, we measured the fold-change in the concentration of 18 phosphoproteins, in young and senescent cells, upon stimulation with 6 cytokines and constructed a signaling network for both populations simultaneously. The MetaCore database was used to assemble the pool of possible interactions.

The responses for young and senescent fibroblasts were compared and we observed significantly higher overall responsiveness of proliferative signals in young cells and potential insulin resistance in senescent cells. Multiple signals associated with the Senescence Associated Secretory Phenotype (SASP) are activated in senescent cells, mainly in response to *EGF, IL1A* and *TNF* stimulation. *SASP* involves the secretion of inflammatory cytokines, growth factors, and proteases and is a characteristic feature of senescent cells. SASP has been associated with many age-related diseases, including type 2 diabetes and atherosclerosis. In our model, the *NFKB* pathway, which is highly involved in the establishment of *SASP* is activated through downregulation of *NFKBIA* by upstream signals and upregulation of *RelA/p65* [22]. The activation of *ERK1/2* is also known to affect the secretion of interleukin 6, the most prominent cytokine of the *SASP* [30] and *MYD88* activation through the interleukin receptor has been associated with *SASP* as well [29].

Another highly studied pathway in aging is the insulin and *IGF1* signaling (IIS) pathway. The attenuation of *IIS* has been shown to promote longevity and extend the lifespan of various organisms [26]. However, *IIS* is also known to decline during normal and accelerated aging [27]. The complexity of the IIS pathway is further increased by evidence that hybrid INSR-IGF1R complexes are formed and have distinct affinities for insulin, IGF-1 and IGF-2 [31]. In this model, we propose that the attenuation of the *IIS* response in senescent cells is mediated through lower responsiveness of *INSR* but not *IGF1R.* While *IGF1R* and *INS’R* share common downstream pathways, their biological activity is not identical. *INSR* mainly controls glusose uptake and metabolism, while *IGF1R* plays a proliferative and anti-apoptotic role [20].

In conclusion, this paper introduced a new logic-based framework for constructing biological networks that allow researchers to model more complex experiments as well as dependencies across networks. To demonstrate its capabilities, we modeled the signaling network of HFL-1 cells as they age. The results presented are preliminary and in the future, we plan to model more types of “-omics” data and time-points in our model.

## Supporting information

**S1 Supporting Information Data Preprocessing.** Data preprocessing pipeline to clean and reformat the Luminex Xmap data in order to run the analysis.

**S2 File luminex_measurements.** Primary data used for the analysis in long format.

**S3 File code.** Code to run the analysis.

**S4 File solutions.** spreadsheet of the results.

## Acknowledgments

This work was supported by the following Research Funding Programs: TS and LGA were supported by ERC “Investing in knowledge society through the European Social Fund” (Grant no. ERC-11/MIS:374071). NC, AC and LGA were supported by Thales “MAESTRO” (Grant no. MIS:377001). Both programs were co-financed by the European Union through the European Social Fund (ESF) and Greek national funds through the Operational Program “Education and Lifelong Learning” of the National Strategic Reference Framework (NSRF).

IB’s PhD thesis is supported by a scholarship from the State Scholarship Foundation in Greece (IKY) (Operational Program “Human Resources Development - Education and Lifelong Learning”, Partnership Agreement (PA) 2014-2020).

